# Entire genome transcription across evolutionary time exposes non-coding DNA to *de novo* gene emergence

**DOI:** 10.1101/017152

**Authors:** Rafik Neme, Diethard Tautz

## Abstract

Even in the best studied Mammalian genomes, less than 5% of the total genome length is annotated as exonic. However, deep sequencing analysis in humans has shown that around 40% of the genome may be covered by poly-adenylated non-coding transcripts occurring at low levels^1^. Their functional significance is unclear^2,3^, and there has been a dispute whether they should be considered as noise of the transcriptional machinery^4,5^. We propose that if such transcripts show some evolutionary stability they will serve as substrates for *de novo* gene evolution, i.e. gene emergence out of non-coding DNA^6–8^. Here, we characterize the phylogenetic turnover of low-level poly-adenylated transcripts in a comprehensive sampling of populations, sub-species and species of the genus *Mus*, spanning a phylogenetic distance of about 10 Myr. We find evidence for more evolutionary stable gains of transcription than losses among closely related taxa, balanced by a loss of older transcripts across the whole phylogeny. We show that adding taxa increases the genomic transcript coverage and that no major transcript-free islands exist over time. This suggests that the entire genome can be transcribed into polyadenylated RNA when viewed at an evolutionary time scale. Thus, any part of the “non-coding” genome can become subject to evolutionary functionalization via *de novo* gene evolution.

---

Genes can emerge *de novo* from non-genic regions of the genome^9–11^. Newly arising transcripts are initially usually non-coding, can later acquire functional open reading frames^12–14^ and can quickly become essential^15^. A number of possibilities have been discussed by which new transcripts can arise, including single mutational events^10^, stabilization of bi-directional transcription^16^ and insertion of transposable elements with promotor activity^17^. These events were initially thought to be rare^6^, but an increasing number of studies show that *de novo* gene emergence is a rather active mechanism^18–22^. Surveys across phylogenetic times have shown that the highest gene emergence rates are found in youngest taxa^7^. This led to the prediction that high emergence rates must be balanced with high loss rates, because gene numbers do not grow much over time^7^. A comparison of open reading frame turnover rates of *de novo* evolved genes among *Drosophila* species has shown that this is indeed the case^22^.

Here we assessed the numbers of new transcript gains in a comprehensive phylogenetic framework. Given that the emergence of a new stable transcript is a prerequisite for evolving a new functional gene, we expect that the transcript emergence rate is a key parameter in the process of *de novo* gene emergence. Stable transcripts can be created out of combinations of cryptic functional sites, including a minimal promoter, splicing signals and poly-adenylation sites. This applies to completely new transcripts and to modification of existing transcripts by adding new exons from non-coding DNA. Given the widespread presence of cryptic functional sites in genome sequences, a single mutational step can convert a non-transcribed or a spuriously transcribed genome region into a stable transcript, as has been shown for *Pldi,* the first documented *de novo* gene in the mouse^10^.

Importantly, once a genomic region becomes transcribed, most subsequent mutations within the transcribed region will not lead to a loss of the transcript, since only a few sites are responsible for active and stable transcription. Hence, one can predict that the *de novo* transcript emergence dynamics would show a higher rate of gain than loss at short evolutionary time scales. Hence, transcriptional gain would constitute a powerful mechanism to continuously expose new genome regions to evolutionary testing, providing the fuel for *de novo* gene emergence.

To test this prediction, we selected species, subspecies and populations related to the house mouse (*Mus musculus* - suppl. Table S1) as a phylogenetic framework for identifying the emergence and loss of new transcripts. The taxa chosen span approximately 10.5 Myr of evolutionary divergence and represent up to ∼5.6% overall genomic divergence (Figure 1, suppl. Table S2). Using such closely related taxa ensures that most neutrally evolving sequences can be reliably mapped across all species^23^. We generated genome sequences for species without published genomes, and transcriptome sequences for brain, liver and testis for all taxa (suppl. Tables S3-S5).

**Figure 1.**
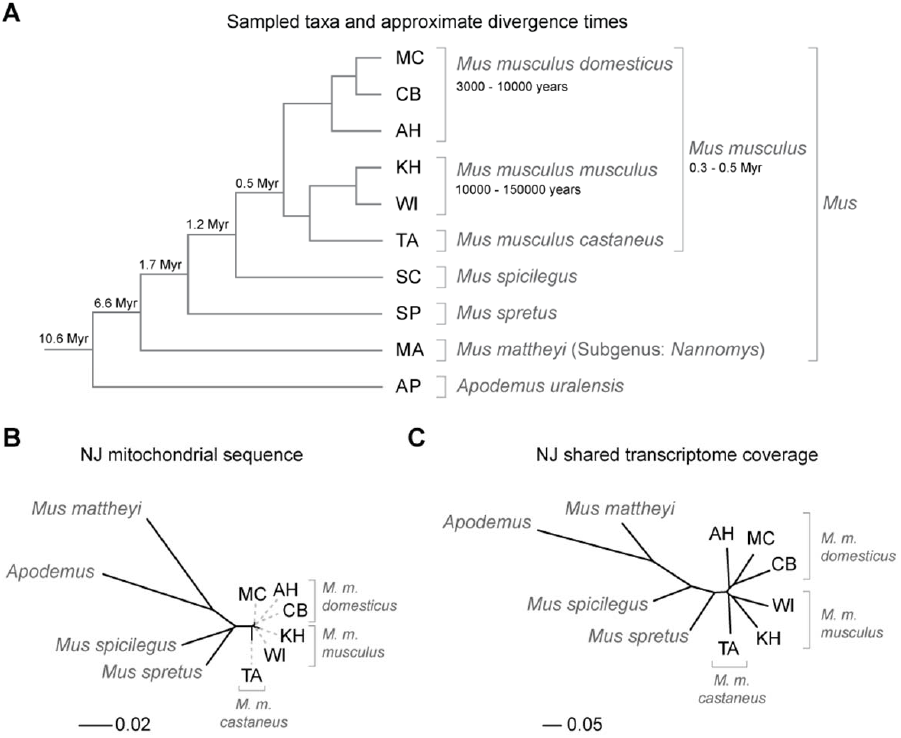
(A) Schematic relationships and approximate divergence times (see Methods) of the taxa under study. Tree branches are not shown to scale. (B) Molecular phylogeny based on whole mitochondrial genome sequences as a measure of molecular divergence (black lines represent the branch lengths, dashed lines serve to highlight short branches). (C) Tree based on shared transcriptome coverage of the genome. The percentage of shared transcripts mirrors the phylogenetic relationships between the studied taxa. MC: *M. m. domesticus* from France. CB: *M. m. domesticus* from Germany. AH: *M. m. domesticus* from Iran. KH: *M. m. musculus* from Kazakhstan. WI: *M. m. musculus* from Austria. TA: *M. m. castaneus* from Taiwan.

For comparative transcriptome analysis, we identified all mappable regions of the *M. m. domesticus* reference genome (C57Bl/6)^24^ using genomic reads from all studied species. We call this the “common genome”, representing the total genome where transcript mapping across taxa is reliable. We used a mapping algorithm that was specifically designed to deal with the polymorphisms occurring under cross-species mapping conditions^23^.

We first focused on genome-wide signals of transcriptional activity to identify the origin of new transcripts within the phylogeny (suppl. Table S8). For this purpose, we determined the base-wise transcriptome coverage from poly-adenylated RNA for each species. This measure of coverage includes both annotated genes and previously un-annotated transcripts, whereby the latter are the majority. We set single read coverage as the lower cutoff because we were specifically interested in detecting low-level transcription as an early sign of *de novo* gene emergence. However, we also report results using a stringent cutoff of five reads for comparison (the median coverage across all transcripts is 3.7).

When comparing transcriptome coverage among taxa, we find that the overall proportion of shared transcripts is higher for closely related taxa than for distantly related pairs. Consequently, a phylogenetic tree reconstructed based on shared transcript coverage mirrors the species tree (Figure 1B, C). This detectable phylogenetic signal in transcription coverage suggests that transcripts gained at a given point in evolutionary time are sufficiently stable to be retained in sister taxa, implying that they can become exposed to evolutionary testing.

The total transcript coverage of the common genome across all species combined for all three tissues is 67% in our data set. Coverage was highest in testis (53.4%), intermediate in brain (41.5%) and lowest in liver (23.5%) (Figure 2A). When comparing all transcriptional gains versus losses across the surveyed phylogeny, we observe that gains are indeed more frequent than losses (Figure 2B, C), thus confirming our initial hypothesis.

**Figure 2.**
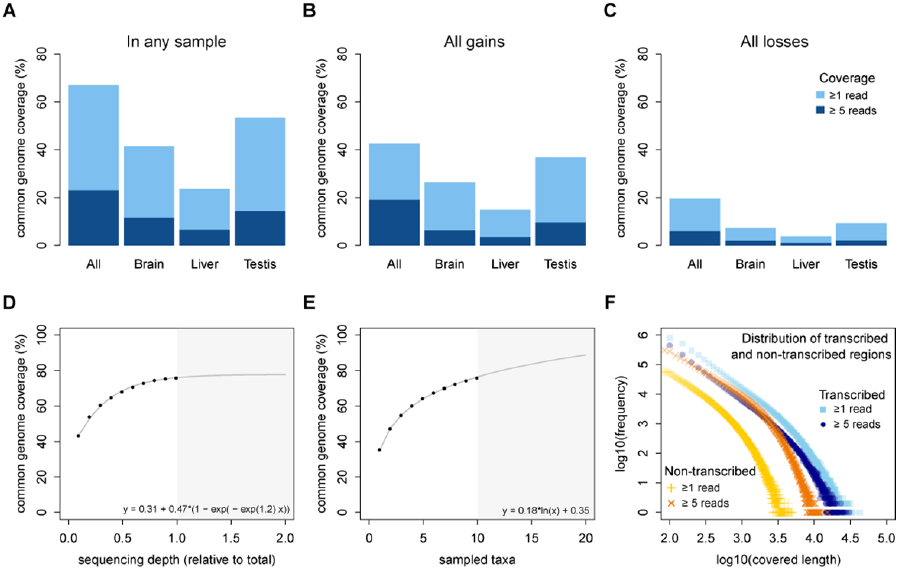
Coverage and phylogenetic turnover of base-wise transcription of the common genome. (A) Coverage across all taxa based on similar sequencing depth for each tissue (suppl. Table xx). (B, C) All gains (B) and all losses (C) along the phylogeny, assuming maximum parsimony (equal gain and loss probability). Light blue represents regions with base-wise coverage of least one uniquely mapping read, dark blue represents regions of base-wise coverage of at least five uniquely mapping reads. (D, E) Rarefaction, subsampling and saturation patterns using all available samples, including deeper sequencing of the brain samples. (D) sequencing depth saturation as estimated from a non-linear regression with asymptotic behavior, (E) sequencing depth saturation as estimated from an increase in the number of taxa. Black dots indicate increases per sub-sampled sequence fraction or taxon added from our dataset. Gray line indicates the predicted behavior from the indicated regression, and gray area shows the prediction after doubling the current sampling either in sequencing effort (D) or additional taxa (E). Each analysis was tested for logarithmic and asymptotic models and best fit was selected by AIC and BIC (see Methods). Standard deviations are too small to become visible in the plots. (F) Comparative analysis of lengths of regions transcribed or not transcribed across all data (including deeper brain sequencing) in all samples. Size distribution of regions not covered in any transcript (yellow) or with less of five transcripts (orange) compared to size distribution of regions with at least one transcript (light blue) or at least five transcripts (dark blue).

Recording base-wise coverage without paying attention to gene models entails the risk that one is also measuring transcriptomic noise, i.e. spurious random transcriptional initiation in a subset of cells of the tissue under investigation. We have specifically explored the noise issue through deeper sequencing of the brain samples of all taxa. The brain is a complex tissue in which some transcripts are expected to occur only in a small subset of cells. These rare transcripts should become detectable by deep sequencing and thus transcript coverage should increase with more reads available. This is indeed the case; however rarefaction curves and their projections reach saturation for each of the taxa (suppl. Table S6 and Figure S1). This observation argues against a significant amount of transcriptional noise in our data, since noise should lead to a continuous increase of coverage with sequencing depth, at least if noise is randomly distributed across the genome and across the cells of the tissue. Further, it rules out a possible problem with DNA contamination, as this would also be expected to rise with increasing sequencing depth.

We have further explored the rarefaction principle to assess whether adding more sequences or more taxa to the data set leads to higher transcriptomic coverage of the common genome. Taking all aggregated reads (including the additional deep sequencing data from brain) across all tissues for all taxa, saturation is reached at 78.5% coverage for sequencing depth (Figure 2D), but no saturation is reached with the number of taxa used here (Figure 2E). Hence, adding more taxa within this phylogenetic framework, for example species and sub-species on the *Apodemus* branch, would predictably lead to increasingly higher transcriptomic coverage of the common genome, up to the entire genome when ∼38 taxa were surveyed within the phylogeny (based on the intercept of the regression curve).

This analysis suggests that there may be no regions that are not transcribed at some point in phylogenetic time. However, genome annotations in a given species usually show an uneven distribution of transcripts; some regions harbor many clustered transcripts and other regions are nearly devoid of transcripts (“gene deserts”). We compared regions devoid of any transcriptomic coverage in our taxonomic sample to regions that show transcription in at least one sample. We find that transcribed regions are more abundant and larger on average than non-transcribed regions (Figure 2E). The maximum length of non-transcribed regions is ∼ 20kb (at ≥1 reads) or ∼50kb (at ≥5 reads), suggesting that large gene-free “deserts” are in principle accessible to transcription and possible regulatory constraints^25^ do not fully prevent their transcription. In fact, the mouse de novo gene *Pldi* has arisen within a gene desert^10^.

Most *de novo* transcripts are expected to be neutral, but some may turn into more stable proto-genes^6,19^ that can eventually become functional, either as regulatory RNA, or by acquiring a functional reading frame. To identify candidate proto-genes we used algorithms that are able to reconstruct transcriptional islands and splice junctions (STAR^26^/cufflinks^27^) and join them into predicted gene models (suppl. Table S9). We did this for each taxon separately and assessed the gain and loss patterns of these transcripts in a phylogenetic context. Excluding all previously annotated transcripts, we find a total of 17,746 new candidate proto-genes, distributed across all taxa (suppl. Figure S2). When looking only at gains of proto-gene transcripts in the terminal branches, we find that about 1,300 new proto-genes are gained per million years (Figure 3A). Interestingly, at least 3,000 proto-genes are already present at the youngest divergence level, implying within-species polymorphism that was also described for *Drosophila*^21^.

**Figure 3.**
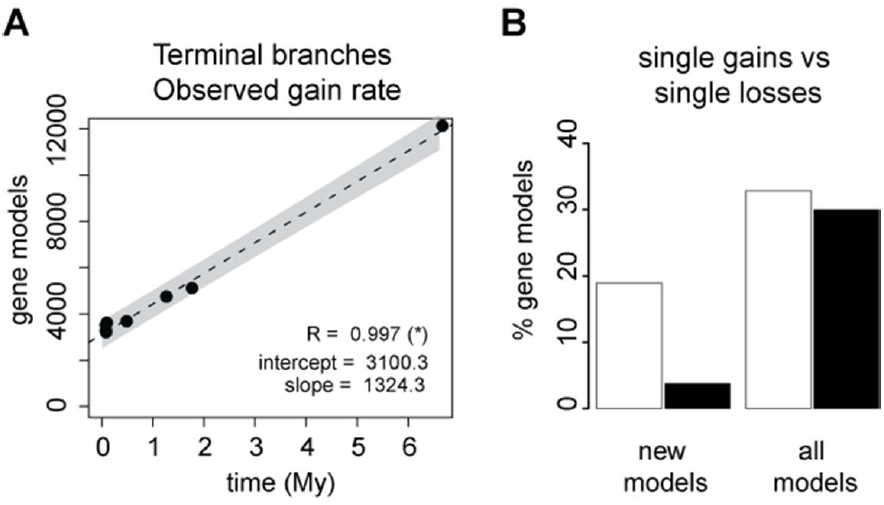
Turnover of proto-gene candidates. (A) Linear regression of observed gains across the phylogeny, using *Apodemus* as outgroup, and counting only gains in terminal branches. Regression coefficient is significant at p < 0.01. The grey area shows the 95% confidence interval of the regression. (B) Single gains versus single losses of a proto-gene candidate, for ‘new’ gene models, i.e. absent in *Apodemus*, and versus all detected models, i.e. including the whole gene set. The ratio of gain to loss of new models is significantly different from the observed for all models (Fisher’s exact test, p < 0.01).

When counting gains versus losses of proto-genes, we find again higher numbers gained than lost over short phylogenetic times. However, gains and losses balance out over longer evolutionary times when including the whole phylogeny and all annotated genes (Figure 3B). This pattern confirms the two essential predictions we made about *de novo* gene emergence: (1) newly acquired transcripts are not easily lost and thus have a life-time sufficient for evolutionary testing and (2) genes do not accumulate over time because gain and loss rates are balanced across longer time spans. Hence, when a given taxonomic lineage gains many *de novo* genes, it will lose some of its older genes.

Our analysis is conservative in several respects. First, we focused on poly-adenylated transcripts, thereby avoiding inclusion of RNA fragments that have been processed (i.e. excised introns) and randomly transcribed fragments. However, this means we also exclude RNAs which are not transcribed by RNA polymerase II, such as tRNAs, snRNAs and ribosomal RNAs. The human ENCODE data suggest that such non-poly-adenylated RNAs are abundant^1^ and it is likely that proto-genes can arise from those transcripts as well, i.e. we are likely underestimating the proto-gene emergence rate. Second, we focused on three tissues and one developmental stage only. Although we included testis and brain, which are known to have the highest diversity of transcripts^28^ we can also expect that including more tissues and developmental stages would further increase the transcriptomic coverage. Taking these factors into account, as well as the fact that increased taxonomic representation shows no signs of saturation with respect to transcriptomic coverage (Figure 2E), it seems reasonable to conclude that when measured at a phylogenetic time scale, the entire genome can become subject to transcription.

Pervasive transcription of the genome was noted soon after deep sequencing approaches became possible and this pattern was systematically explored in the ENCODE projects^1,29^. While the functional significance of pervasive transcription is a matter of continuous dispute^4,5,30^, our results provide an evolutionary dynamics perspective on this question where emergence, functionalization and decay of gene functions should be seen as an evolutionary life cycle of genes^8^. *De novo* gene birth should no longer be considered as the result of unlikely circumstances, but rather a mechanism of testing genome regions for their adaptive potential. Within this evolutionary perspective, any part of the genome – “junk” DNA included – has the possibility to become useful.

## Supplementary information

Supplementary_files includes Tables S1-S7 and Figures S1 and S2. Supplementary_TableS8 - Excel file with gain loss patterns of transcript coverage per branch of the phylogeny. Supplementary_TableS9 - Excel file with list of all gene models and gain/loss pattern along branches.

## Acknowledgements

We thank the C. Pfeifle and the mouse team for providing the animals, J. Altmüller and C. Becker for sequencing, B. Harr, A, Nolte, L. Pallares and L. Turner for comments on the manuscript and the members of our group for discussions and suggestions. Special thanks to F. Sedlazeck for bioinformatic advice and provision of software before publication. R.N. was supported by a PhD fellowship of the IMPRS for Evolutionary Biology during the initial phase of the project. The project was financed through an ERC advanced grant to D.T. (NewGenes - 322564).

### Author contribution

DT and RN conceived the project, RN did the experimental work and data analysis, DT and RN discussed the data interpretation and wrote the manuscript.

## Methods online

### Origin of the sampled taxa

We selected ten taxa, ranging from population level through sister genera (Figure 1A).

The youngest divergence point sampled, at about 3,000 years, corresponds to the split between two European populations of *Mus musculus domesticus* ^1^ one from France (Massif Central = MC) and one from Germany (Cologne-Bonn area = CB) ^2^. These European populations in turn have diverged from an ancestral *M. m. domesticus* population in Iran (Ahvaz = AH) about 12,000 years ago ^1^. The European *M. m. domesticus* are also the closest relatives of the reference genome, the C57BL/6J strain ^3^.

We included two populations of *Mus musculus musculus*; one from Austria (Vienna = WI) and one from Kazakhstan (Almaty = KH). These two populations are supposed to have a longer divergence between then the European *M. m. domesticus* populations, but a more accurate estimate is currently not available. We set the divergence for analyses at around 10,000 years as an approximate estimate.

*M. m. domesticus* has diverged from *M. m. musculus* and *Mus musculus castaneus* about 0.4 to 0.5 million years ago, with a subsequent divergence, not long after, between *M. m. musculus* and *M. m. castaneus* ^4^. We included *M. m. castaneus* from Taiwan as a representative of the subspecies.

To account for longer divergence times, we included *Mus spicilegus* (estimated divergence of 1.2 million years); *Mus spretus* (estimated divergence of 1.7 million years) ^4^; *Mus matteyii* (subgenus *Nannomys*), the North African miniature mouse (estimated divergence of 6.6 million years) ^5,6^, and *Apodemus uralensis*, the ural field mouse (estimated divergence of 10.6 million years) ^6^.

The population-level samples (*M. m. domesticus* and *M. m. musculus*) included are maintained under outbreeding schemes, which allows for natural polymorphisms to be present in the samples. All other non-population samples are kept as more or less inbred stock, and therefore fewer polymorphisms are expected. All mice were obtained from the mouse collection at the Max Planck Institute for Evolutionary Biology, following standard rearing techniques which ensure a homogeneous environment for all animals. Mice were maintained and handled in accordance to FELASA guidelines and German animal welfare law (Tierschutzgesetz § 11, permit from Veterinäramt Kreis Plön: 1401-144/PLÖ-004697).

A total of 60 mice were sampled, as follows: Eight male individuals from each population-level sample (outbreds), Iran (AH), France (MC), and Germany (CB) of *Mus musculus domesticus*, and Austria (WI) and Kazakhstan (KH) of *Mus musculus musculus*. Four male individuals from the remaining taxa (partially inbred): *Mus musculus castaneus* (TA), *Mus spretus* (SP), *Mus spicilegus* (SC), *Mus mattheyi* (MA) and *Apodemus uralensis* (AP). Mice were sacrificed by CO_2_ asphyxiation followed immediately by cervical dislocation. Mice were dissected and tissues were snapfrozen within 5 minutes post-mortem. The tissues collected were liver (ventral view: front right lobe), both testis and whole brain including brain stem.

### Genome sequencing

One individual from each of *M. spicilegus*, *M. mattheyi*, and *Apodemus uralensis* were selected for genome sequencing. DNA was extracted from liver samples. DNA extraction was performed using a standard salt extraction protocol. Tagged libraries were prepared using the Genomic DNA Sample preparation kit from Illumina, following the manufacturers’ instructions: After library preparation, the three genome samples were pooled together and run in a whole IlluminaHiSeq 2000 flow cell (8 lanes, approximately 2.6 lanes per sample). Library preparation and sequencing was performed at the Cologne Center for Genomics.

The genome from the strain SPRET/EiJ derived from *Mus spretus* was taken from ^7,8^, and was downloaded from the European Nucleotide Archive (ENA) - accessions ERS076388 and ERS138732.

### Transcriptome sequencing

The sampled tissues of each taxon were used for RNA extraction with the RNAeasy Mini Kit (QUIAGEN) and pooled at equimolar concentrations. RNA quality was measured with the Agilent RNA Nano Kit, for the individual samples and pools. Samples with RIN values above 7.5 were used for sequencing. Library preparation was done using the Illumina TruSeq library preparation, with mRNA purification (PolyA selection), following manufacturers’ instructions. Sequencing was done in Illumina HiSeq 2000 sequencer. Libraries for each group were tagged, pooled and sequenced in a single lane, corresponding to approximately one third of a HiSeq2000 lane. Additional sequencing of the brain samples was performed to identify potential limitations in depth of sequencing. For this, each brain library was sequenced on a full lane of a HiSeq2000. All library preparation and sequencing was done at the Cologne Center for Genomics (CCG).

### Raw data processing

All raw data files were trimmed for adaptors and quality using Trimmomatic ^9^. The quality trimming was performed basewise, removing bases below quality score of 20 (Q20), and keeping reads whose average quality was of at least Q30. Reads whose trimmed length was shorter than 60 bases were excluded from further analyses, and pairs missing one member because of poor quality were also removed from any further analyses.

### Mapping

The reconstruction of transcriptomes using high-throughput sequencing data is not trivial when comparing information across different species to a single reference genome. This is due to the fact that most of the tools designed for such tasks do not work in a phylogenetically aware context. For this reason, any approximation which deals with fractional data (i.e. any high-throughput sequencing setup available to this date) is limited by the detection abilities of the software of choice and by the quality of the reference (transcriptome and genome).

Given the high quality state of the mouse genome repositories, we decided to take a reference-based approach, in which all analyses are centered in the reference genome of the C57BL/6 laboratory strain of *Mus musculus domesticus*. This enables direct comparisons across all species, with an obvious cost introduced by the mapping of distantly related genomes.

For general comparisons, transcriptome and genome sequencing reads were aligned against the mm10 version of the mouse reference genome from UCSC ^10^ using NextGenMap which performs extremely well with divergences of over 10% compared to other standard mapping software ^11^. The program was run under default settings, except for --strata 1 and --silent-clip. The first option enforces uniquely mapping reads and the second drops the unmapped portion of the reads, to avoid inflating coverage statistics. This is particularly relevant around exonintron boundaries, where exonic reads are forced into intronic regions unless this option is set.

Genomic reads were used to as empiric mapability, i.e. to identify which regions can be reliably detected. We limited our analyses to regions in the reference genome which could be mapped at least 5 times from genomic reads from all other species (5x coverage). This is the portion we call the ‘common genome’ in downstream analyses. It is important to highlight that this is not the same as synteny, since we did not perform any co-linearity analyses between fragments, but rather represents the mere presence in the species, in any possible order.

Furthermore having genomic reads enables the detection of true absences in transcriptional activity from absences of genome regions, which would show similar patterns in transcriptome-only analyses.

### Reconstruction of gene models

Due to the fact that NextGenMap is unable to perform split read analyses we opted for more standard tools to reconstruct gene models from the data. For this we used STAR ^12^ to map reads to the reference, followed by cufflinks ^13^ to obtain automated gene and transcript annotations for each species. The annotation file contains models for expressed transcripts with splicing information (exon annotation) when available. All annotations were merged using cuffmerge to generate a final annotation that includes gene models present at least once in the total sample. Mono-exonic models shorter than 500 bases or contained within introns of multi-exonic transcripts were excluded from analyses.

### Parsimony gain and loss mapping

We estimated gain and loss events given the phylogenetic distribution of presence and absence of transcription at a given position or for a given gene model using maximum parsimony (based on GLOOME, ^14^, the assumption that gains and losses are equally likely, and a fixed tree describing the relationships between taxa.

### Genome-wide estimation of transcriptional gains and losses

Genome-wide estimates of gain and loss of transcription were done at the nucleotide level, considering only regions within the common genome.

Normalized versions of each set of aligned reads were generated by subsampling (samtools view –s x ^15^; where x is the proportion of each individual sample that matches the least abundant sample). Normalization was done only across tissues and not between them. Normalized samples were merged at the species level to obtain a species-wide transcription sample. Aligned reads in BAM format were converted to BedGraph format for phylogenetic comparisons of format using bedtools ^16^. Two parallel sets of comparisons were made: i) using all coverage information from uniquely mapping reads, thus representing ‘absolute’ (minus normalization) coverage, and ii) using a threshold of at least 5 uniquely mapped reads, thus representing ‘stringent’ coverage. Coverage files were compared between groups using multiIntersectBed ^16^, to obtain the portions shared between all possible combinations.

Each possible combination can be also interpreted as a binary presence/absence pattern. We summarized the total amount of nucleotides in each specific pattern. Each pattern received a fixed amount of gains and losses, consistent with the parsimony assumptions using GLOOME ^14^ in maximum parsimony mode. For example, the pattern that indicates presence in German *M. m. domesticus* (CB), but absence in all other groups, corresponds to one very recent gain and zero losses. The pattern that indicates presence in German and Iranian *M. m. domesticus* (CB and AH), but absence in all other groups, corresponds to one gain (ancestor of *M. m. domesticus*) and one loss (after divergence of French *M. m. domesticus*). In this context, we identify monophyletic gains as stable, i.e. transcription is present in all derived groups after estimated gain, and unstable, i.e. transcription lost in at least one derived group after gain.

### Gene gain and loss rates from gene models

Gene models derived from STAR alignments and cufflinks reconstructions were used to calculate the rate at which gene-like entities are gained or lost along the phylogeny. A single unified annotation for all species was generated with cuffmerge (see “Gene model reconstruction” above) and FPKM values were obtained from mapped reads for each tissue and species. FPKM of 0.1 was set as a threshold to define the presence or absence of a gene model in a given sample. Similar to the reconstruction of genome-wide patterns, a maximum parsimony framework was employed, assuming that gains and losses have equal probabilities of occurrence.

Rates of gain and loss of gene models were estimated from linear regressions between time and gain or loss using R, with the lm() function from the stats package ^17^. Rates were calculated for each tissue and each possible combination of tissues (a gene can be differentially present or absent in a species), to obtain tissue-specific gains and losses of models.

### Reconstruction of phylogenetic relationships

Using a manhattan distance matrix from the summarized transcriptional coverage (suppl. Table S8), we constructed a neighbor-joining (NJ) tree that describes the proximity between any two taxa based on the shared transcriptomic coverage between them. In this representation, closely related organisms have more shared transcriptomic coverage than distantly related organisms. Analyses were performed in R, using the function dist() from the stats package and nj() from the ape package ^18^.

Additionally, whole mitochondrial genomes were obtained for each taxon as consensus sequences from mapped reads using samtools mpileup ^15^. The sequences were aligned with MUSCLE ^19^, and a NJ tree was constructed with the dist.dna() and nj() functions from the ape package ^18^. All NJ trees were tested with 1000 bootstraps with the boot.phylo() function from the ape package ^18^. Reported trees have a support of 60% or greater.

### Rarefaction and subsampling

Transcriptome experiments tend to be limited by the depth of sequencing, with highly expressed genes being relatively easy to sample, and rare transcripts becoming increasingly difficult to find. Given the large amount of data generated, we investigated if our data shows signals of coverage saturation from subsets of the data of different sizes. The total experiment, comprising ten taxa, corresponds to 6.4 x 10^9^ reads (or 6.4 billion reads). We subsampled (samtools view -s) portions of mapped reads for each taxon, ranging between 10% to 90%, at 10% intervals. The observation of coverage saturation in this case would indicate that our sequencing efforts likely cover most of the transcribed regions of the common genome.

In parallel, we estimated the individual and combined contribution of each taxon to the transcriptomic coverage of the common genome. Not all samples have the same phylogenetic distance to each other (some species have more representatives than others). To account for this we generated one hundred arrays of the ten taxa with random order, and recorded the coverage after the addition of each taxon in each array. The observation of coverage saturation in this setup would indicate that taxonomic sampling is sufficient to cover most of the potentially transcribed regions of the common genome. In order to estimate whether our data continued to increase or approached saturation, we tested two alternative models: a generalized linear model with logarithmic behavior (ever increasing) or a self-starting nonlinear regression model (saturating). Best fit was decided based on the lowest AIC and BIC values. Analyses were performed in R, using the functions glm(), nls(), SSasymp(), BIC() and AIC() from the stats package ^17^.

### Analysis of transcribed and non-transcribed regions across the genome

Transcribed and non-transcribed regions larger than 100 nucleotides were defined by the continuous presence or absence of transcriptomic coverage from mapping information of each taxon and tissue. Combined transcribed regions across species were obtained as mentioned before, and combined non-transcribed regions across species were generated by subtracting transcribed regions from the common genome.

